# TranSiGen: Deep representation learning of chemical-induced transcriptional profile

**DOI:** 10.1101/2023.11.12.566777

**Authors:** Xiaochu Tong, Ning Qu, Xiangtai Kong, Shengkun Ni, Kun Wang, Lehan Zhang, Yiming Wen, Sulin Zhang, Xutong Li, Mingyue Zheng

## Abstract

With the advancement of high-throughput RNA sequencing technologies, the use of chemical-induced transcriptional profiling has greatly increased in biomedical research. However, the usefulness of transcriptomics data is limited by inherent random noise and technical artefacts that may cause systematical biases. These limitations make it challenging to identify the true signal of perturbation and extract knowledge from the data. In this study, we propose a deep generative model called Transcriptional Signatures Generator (TranSiGen), which aims to denoise and reconstruct transcriptional profiles through self-supervised representation learning.TranSiGen uses cell basal gene expression and compound molecular structure representation to infer the chemical-induced transcriptional profile. Results demonstrate the effectiveness of TranSiGen in learning and predicting differential expression genes. The representation derived from TranSiGen can also serve as an alternative phenotype information, with applications in ligand-based virtual screening, drug response prediction, and phenotype-based drug repurposing. We envisage that integrating TranSiGen into the drug discovery and mechanism research pipeline will promote the development of biomedicine.

## Introduction

Transcriptional profiling is a powerful tool that is widely used to characterize the phenotype of various cells and organisms. It captures the landscape of thousands of genes in different biological circumstance at a holistic level, providing abundant information on cellular and organismal status. Transcriptomics data analysis helps us further understand the changes in cellular functionality during the pathogenesis of disease and identify the regulatory mechanisms of cells in response to different perturbations.

With the significant advancements in high-throughput RNA sequencing (RNA-seq) technologies, the availability of comprehensive and systematic perturbational gene expression profiles has greatly expanded. Connectivity Map (CMap)^1^, a public database of perturbational profiles, initially contained data of 164 different bioactive small molecules in four cell lines. To obtain large-scale CMap data in a cost-effective and high-throughput manner, a novel gene expression profiling platform called L1000 was introduced by the Library of Integrated Network-based Cell-Signature (LINCS) program. L1000 measures the expression of 978 landmark genes to capture most of the information in the full transcriptome^2^. In 2017, the CMap-L1000v1 dataset (Phase I, GEO: GSE92742) was reported to contain L1000 profiles from 42,080 genetic and small-molecule perturbations across approximately 80 cell lines, with ongoing updates. The PANACEA data, being developed by the Columbia Cancer Target Discovery and Development Center, is expected to contain dose-responses and RNA-seq profiles from perturbations with approximately 400 clinical cancer drugs across 25 cell lines^3^. In addition, there are ongoing efforts to collect and organize gene expression profiles from publicly available databases to create standardized and unified transcriptional data. ARCHS4 is a web resource containing RNA-seq data from human and mouse^4^, while ChemPert focuses on RNA-seq data for non-cancer cell lines^5^.

These large-scale perturbational gene expression profiles play an indispensable role in drug discovery and mechanism research. The CMap project proposed a pattern-matching strategy to identify compounds that share a mechanism of action (MOA). This approach led to the discovery of a novel mechanism for gedunin, which acts as an inhibitor of HSP90 function^1^. Associating chemical-induced transcriptional profiles with diseases has several benefits, such as screening candidate compounds for diseases and revealing mechanisms of drug resistance. Chen et al. proposed a method to measure the potency of reversing disease gene expression and quantify the reversal relationship between disease and drug gene expression profiles. Through their systematic exploration, they validated that pyrvinium pamoate, a FDA-approved drug for the treatment of pinworms, also has in vivo efficacy for liver cancer^6^. In addition, Wei et al. identified that rapamycin can reverse glucocorticoid resistance by screening chemical-induced profiles. They discovered that MCL1 is an important regulator of glucocorticoid-induced apoptosis^7^.

Machine learning algorithms have been used to automatically capture relationships between different perturbational profiles. Pabon et al. employed a random forest (RF) model to learn the relationship between chemical-induced profiles and gene knockdown-induced profiles, thereby identifying potential targets of compounds^8^. We proposed a graph convolutional network model SSGCN, which integrates protein-protein interaction networks and detects intricate relationships behind perturbational gene expression profiles. This approach facilitates the inference of protein targets for compounds^9^.

While numerous drug-like molecules have undergone high-throughput transcriptional perturbation experiments, it is not feasible to explore all possible perturbation-cell combinations due to the vast size of the combinatorial space. Therefore, a promising approach is to develop deep learning models capable of learning from high-dimensional data extracted from publicly accessible transcriptional profiles. By utilizing these models to predict gene expression profiles, it becomes possible to access the unexplored space of perturbation transcriptomics. Recently, several studies have explored the use of deep learning models to predict transcriptional profiles for new chemicals. Zhu et al. proposed a deep neural network DLEPS, which is capable of fitting chemical-induced gene expression values without considering the types of cell lines. This model has been utilized to identify potential candidates for obesity, hyperuricemia and nonalcoholic steatohepatitis^10^. On the other hand, DeepCE^11^ and CIGER^12^ leveraged one-hot encoding to discriminate different cell types, and learned from perturbation profiles of various cells. These two models focused on predicting differential gene expression profiles disrupted by novel chemicals, and have been applied to the drug repurposing pipeline using drugs from the DrugBank database^11,12^.

However, existing models that directly fit gene expression values via supervised learning may not effectively distinguish among the true perturbation signals, confounding factors, and high level of noises in expression profiles. Recent studies have explored the use of variational autoencoder (VAE) and its variants for denoising, dimensionality reduction, imputing missing values^13^, and extracting meaningful biological signals^14^ from gene expression profiles. While these studies have demonstrated the impressive capabilities of VAE-based models in processing high-dimensional and noisy transcriptomics data, these analyses are limited to the reconstruction of experimental perturbation profiles and could not be extended to novel perturbations.

To address these limitations, we propose a VAE-based framework called Transcriptional Signatures Generator (TranSiGen) for learning and reconstructing the transcriptional profiles. TranSiGen is designed to alleviate inherent limitations of data through self-supervised representation learning and can be used to infer new perturbational profiles. We evaluated the performance of TranSiGen in fitting basal profiles *X*_1_, perturbational profiles *X*_2_, and the corresponding differential expression genes (DEGs). Furthermore, we analyzed TranSiGen-derived DEGs to assess its ability to effectively learn cellular and compound features from perturbation profiles. In addition, we compared the performance of TranSiGen with other baseline models in the task of inferring DEGs. Finally, we demonstrated the effectiveness of TranSiGen-derived representation as a type of phenotype information by applying it to various downstream tasks.

The highlights of TranSiGen are as follows: (1) TranSiGen uses self-supervised representation learning strategy on a VAE-based model to denoise and reconstruct transcriptomics data. This approach yields impressive performance in inferring chemical-induced transcriptional profiles. (2) The perturbational expression profiles obtained by TranSiGen effectively learn cellular and compound features from data, and it can serve as a new, unified and standardized representation for characterizing chemical-induced phenotype information. (3) TranSiGen-derived representation has demonstrated its applications in a range of downstream tasks, including ligand-based virtual screening, drug response prediction, and phenotype-based drug repurposing for disease.

## Results

### The Overview of TranSiGen

TranSiGen is a VAE-based model that simultaneously learns three distributions: basal profiles without perturbation, perturbational profiles, and the mapping relationship between them. It utilizes a self-supervised representation learning strategy to mitigate noise effects in the transcriptional profile and uncover the signal of perturbation.

The transcriptional profiles used in the model are obtained from level3 data of the newly released CMAP LINCS 2020 dataset^2,15^. These profiles consist of 978 measured landmark genes per profile. Specifically, basal profiles (*X*_1_) represent control profiles treated with DMSO, while perturbational profiles (*X*_2_) represent transcriptional profiles treated with compounds. For each plate, the DMSO-treated control profile from the same plate is selected as *X*_1_, forming a paired *X*_1_∼ *X*_2_. The dataset includes 219,650 ∼ pairs for 8,316 compounds across 164 cell lines. Since L1000 assays are typically conducted with three or more biological replicates, there may be multiple *X*_1_∼ *X*_2_ pairs for a perturbation-cell combination in the dataset. To ensure only one *X*_1_∼ *X*_2_ pair per perturbation on each cell line, the repeated and pairs were further processed using the moderated-Z weighted averages algorithm (MODZ). The processed data consists of transcriptional profiles for 8,316 compounds on 164 cell lines, including 78,569 *X*_1_∼ *X*_2_ pairs (Fig. 1a).

**Fig. 1.**
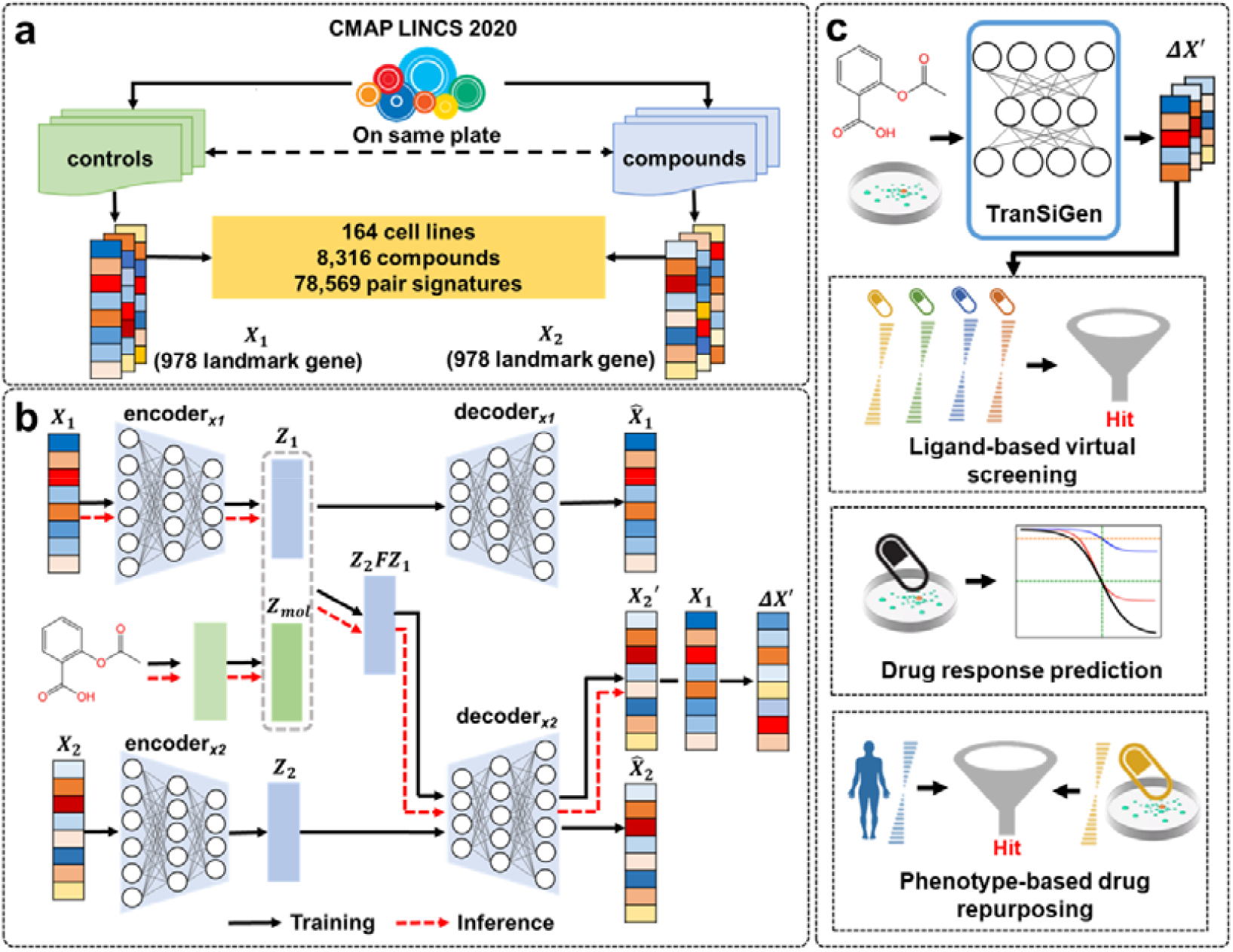
TranSiGen’s architecture and application. **a** The data processing flow for TranSiGen. **b** The architecture and inference process of TranSiGen. **c** The applications of TranSiGen-derived representation.

TranSiGen consists of two VAE models: one encodes basal profiles *X*_1_, and another encodes perturbational profiles *X*_2_ (Fig. 1b and Supplementary Fig. 1). It learns to map from *X*_1_ and the perturbation representation to *X*_2_, which is denoted as 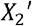. During inference, TranSiGen generates 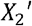 from the input *X*_1_ and notations and the perturbation representation (Fig. 1b), and finally obtains the inferred DEGs 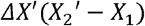 of the compound. A complete list of the symbols used here were summarized in Table 1.

**Table 1.**
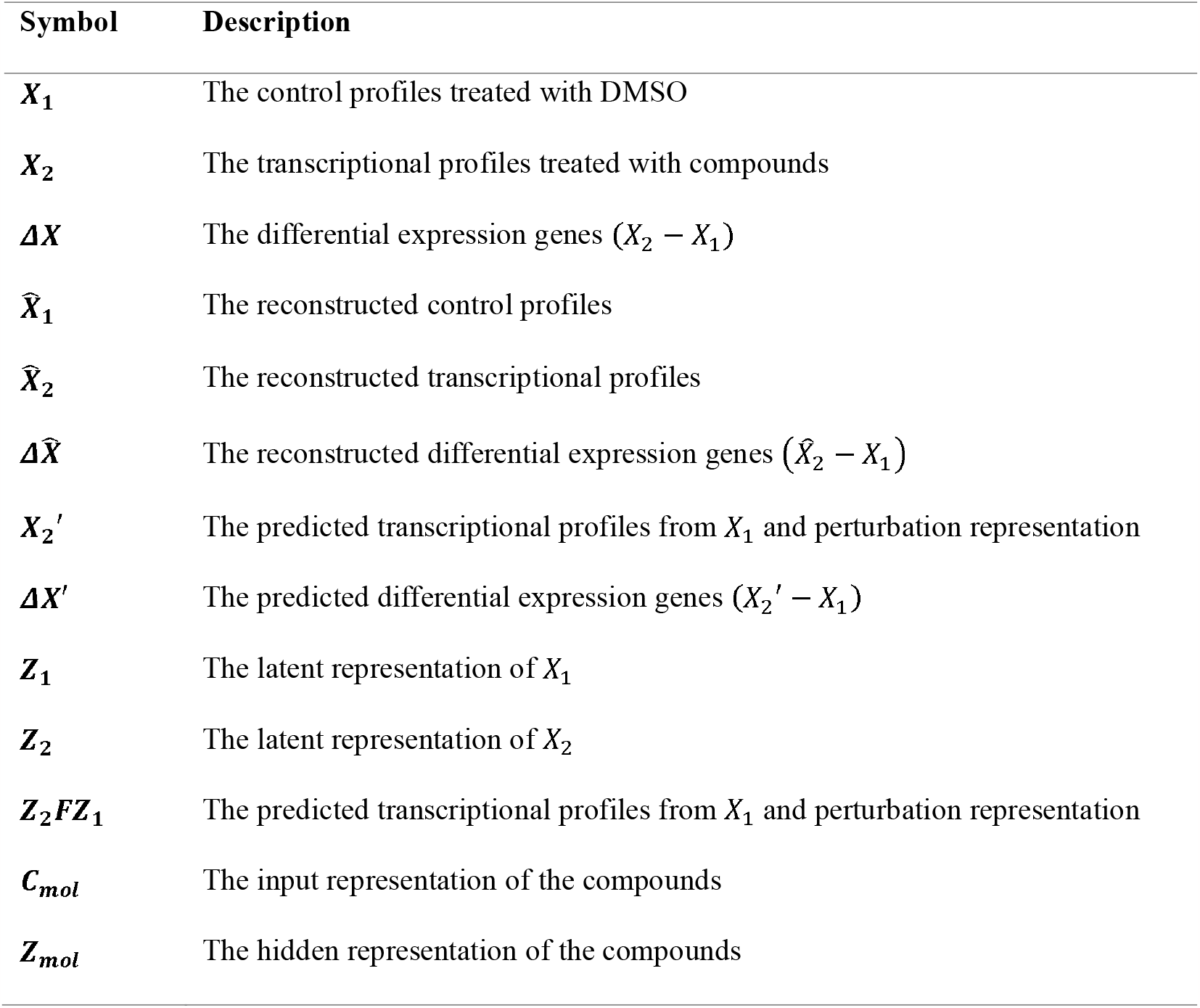
List of symbols and notations used in the paper.

In downstream applications, TranSiGen can generate perturbational profiles for numerous compounds, allowing exploration of a larger space that is not covered by training data. The perturbational representation derived from TranSiGen can be applied to ligand-based virtual screening, drug response prediction in cells, and phenotypic screening of candidate compounds for disease (Fig. 1c).

### TranSiGen enables effective learning for transcriptional profiling

In this study, TranSiGen was used to simultaneously fit the basal profile and the perturbational profile . The model’s performance in learning, and the corresponding DEGs was evaluated individually. As shown in Fig. 2a, TranSiGen exhibits excellent performance in reconstructing and, denoted as and, with the Pearson’s correlation coefficients (PCC) close to 1 (between and, between and). It also performs well in inferring, which is predicted by and compound representation. Compared to directly evaluating the performance of learning and, the corresponding performance in fitting DEGs is slightly decreased, with the PCC 0.734 and 0.619 in reconstructing 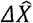 (between 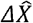 and *ΔX*) and predicting *ΔX*′ (between *ΔX*′ and *ΔX*), respectively. Additionally, the relationship between TranSiGen’s performance and *X*_1_ ∼ *X*_2_ correlation coefficient (R^2^) was analyzed. As shown in Fig. 2b, the sample size of the profiles increases with *X*_1_ ∼ *X*_2_ R^2^, as well as the prediction performance for DEGs. For *X*_1_ ∼ *X*_2_ R^2^>0.8, there is a slight decrease in performance, possibly due to the perturbation effects being too subtle for the model to fully capture. Overall, the model has learned the meaningful mapping from *X*_1_ and compound to *X*_2_.

**Fig. 2.**
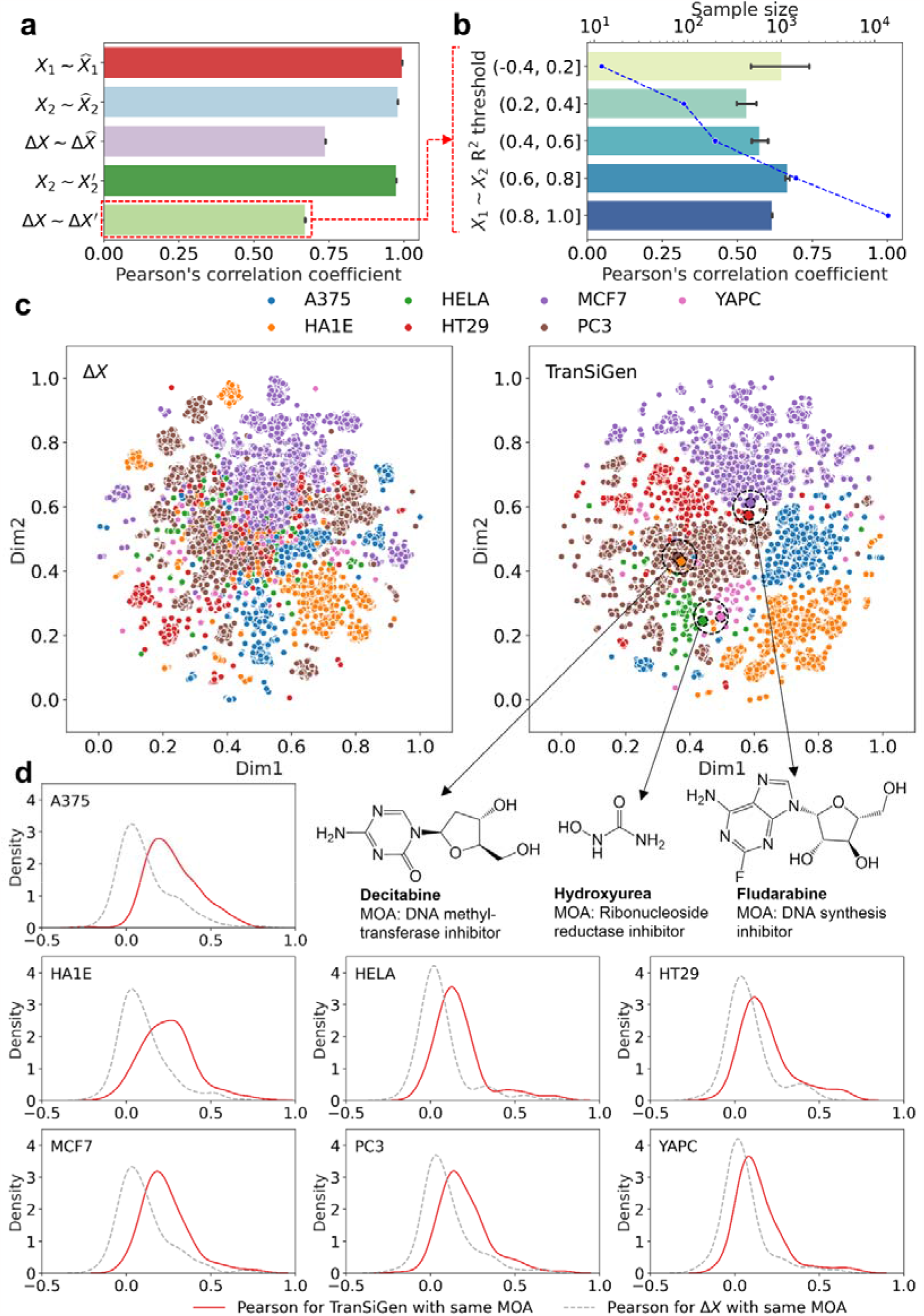
Transcriptional profiling representation learning by TranSiGen. **a** Performance of TranSiGen in transcriptional profiling reconstruction and prediction. **b** The change of TranSiGen’s performance for *ΔX*′ with the correlation between *X*_1_ and *X*_2_. **c** Dimensionality reduction visualization using *ΔX* and TranSiGen-derived *ΔX* ′ for different cell lines. **d** Distribution of Pearson’s correlation coefficients of profiles for the same MOA by *ΔX* and TranSiGen-derived *ΔX*′.

a Furthermore, we evaluated the profiling capabilities of TranSiGen by analyzing its effectiveness in learning cellular and compound representations in *ΔX* ′. Fig. 2c presents a visualization of dimensionality reduction for both experimental *ΔX* and TranSiGen-derived *ΔX* ′, with each point color-coded by cell type. In the case of*ΔX*, there was some clustering of the same cells, but also significant mixing between different cell types. In contrast, TranSiGenderived *ΔX* ′ exhibited clear clustering of the same cells and sharper distinctions between different cell types. This suggests that the representation derived from TranSiGen can more effectively differentiate between various cell types compared to experimental profiling, which is subject to high level of noise. Moreover, for compounds like decitabine, hydroxyurea and fludarabine, the *ΔX*′ for different cells are closely grouped together, indicating their similar perturbation effects across different cells, as they all directly induce cell death due to cytotoxicity^16^. In addition to cytotoxic compounds, we expect other compounds sharing the same MOA to display similar effects on transcriptional profiling. The correlation between compounds with the same MOA was analyzed and is showed in Fig. 2d. TranSiGen-derived representations have higher PCC for compounds with the same MOA than *ΔX*. Meanwhile, when compared to random MOA, TranSiGen-derived representations of the same MOA also exhibit relatively high PCC (Supplementary Fig. 2).

Overall, TranSiGen’s self-supervised representation learning helps denoise and reconstruct transcriptional profiles, effectively identifying and learning meaningful cellular and compound representations from data.

### Comparison with existing models in inferring differential expression genes

In this section, we will compare the performance of TranSiGen with other baseline models in inferring differential expression genes. The models include DLEPS^10^, DeepCE^11^ and CIGER^12^. Among them, DLEPS^10^ predicts one DEG profiling for each compound without considering cell lines, while DeepCE^11^, CIGER^12^, and our model TranSiGen are all capable of inferring cellspecific DEG profiling.

We evaluated the performance of the models using three data splitting methods: random, chemical-blind, and cell-blind. Since not all pairwise combinations of cells and perturbations in the L1000 dataset have experimental profiles, there are missing values in the perturbation-cell combination space. In random splitting (scenario 1), the model predicts DEGs for new combinations of compounds and cells, allowing for the imputation of missing values. In chemical-blind splitting (scenario 2), the model’s capacity to extrapolate profiles for novel compounds (using test compounds not included in the training set) is evaluated. This scenario includes two tests: In scenario 2-1, a dataset with 355 compounds across 7 cells was used to ensure comparability among multiple models. In scenario 2-2, a complete dataset with 8,316 compounds across 164 cells was used to evaluate TranSiGen’ s performance. In cell-blind splitting (scenario 3), cells not included in the training set were used to infer chemical-induced profiles on new cells. TranSiGen characterizes cells using transcriptional profiles without compound treatment, enabling exploration of performance in cell-blind splitting, a scenario that other models cannot address. The model was trained on 10, 50, and 150 cells, and evaluated on 7 new cells. The diagram illustrating the data splitting for chemical-blind and cell-blind scenarios is shown in Fig. 3a.

**Fig. 3.**
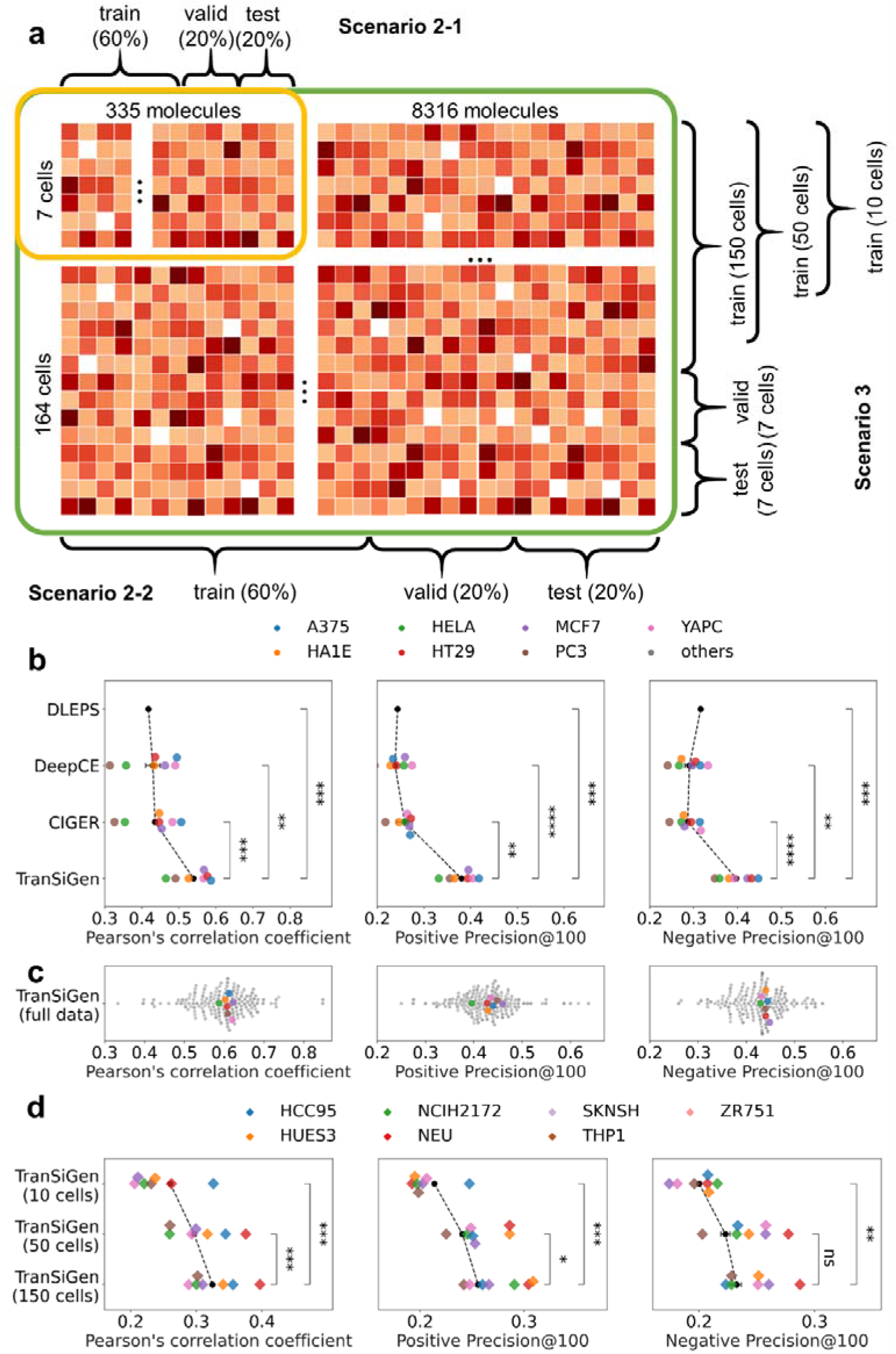
The diagram of data splitting and the performance of inferring DEGs in different scenarios. **a** The diagram of chemical-blind splitting and cell-blind splitting. In scenario 2-1, a dataset of 355 compounds on 7 cell lines is split by compounds, ensuring that test compounds do not seen in the training set. In scenario 2-2, a complete dataset of 8,316 compounds on 164 cell lines is split by compounds. In scenario 3, the complete dataset of 8,316 compounds on 164 cell lines is split by cell lines. The model was trained using the profiling data of 10, 50, and 150 cell lines, and the prediction performance was evaluated on 7 new cell lines. **b** Model performance comparison in chemical-blind splitting. **c** The performance of TranSiGen in chemical-blind splitting by using the full dataset. **d** The performance of TranSiGen in cell-blind splitting by using different numbers of cell lines in the training set. Statistical t-test was applied between the models. (Note: ****, *p <* 0.0001; ***, 0.0001 *< p* ≤ 0.001; **, 0.001 *< p* ≤ 0.01; *, 0.01 *< p* ≤ 0.05 and ns, 0.05 *< p* ≤ 1.0)

According to the results shown in Fig. 3b, TranSiGen performs better than the other three models in scenario 2-1, in terms of the average PCC (represented by the black dots) evaluated on seven cell lines. Additionally, TranSiGen, along with the other two models (DeepCE and CIGER) that consider cell lines contexts, shows similar trends in the prediction performance of DEGs on different cell lines. TranSiGen also outperforms the other three models in terms of Positive Precision@100 and Negative Precision@100 when evaluating metrics focusing on the most significantly up- and down-regulated expressed genes. Furthermore, TranSiGen achieves state-of-the-art results on the full dataset (scenario 2-2), which includes more cell lines and molecules (Fig. 3c and Supplementary Table 1). Compared to the significant performance differences across seven different cells in Fig. 2b, TranSiGen (full data) demonstrates considerable capabilities in inferring DEGs for these seven cell lines by benefiting from more training data. A similar trend is observed in random splitting, where TranSiGen performs better than other models, and it also demonstrates decent performance on the full dataset (Supplementary Table 2). Meanwhile, all models exhibit better performance in random splitting than in chemical-blind splitting due to the inclusion of compounds and cell lines seen during training (Supplementary Table 1 and 2).

**Table 2.** Description of the evaluation metrics.

| Evaluation metric | Equation* |
| --- | --- |
| RMSE | $\sqrt{\frac{1}{n} \sum_{i=1}^n (\Delta X_i - \Delta X'_i)^2}$ |
| Pearson's correlation coefficient | $\frac{cov(\Delta X - \Delta X')}{\sigma_{\Delta X} \sigma_{\Delta X'}}$ |
| Positive Precision @100 | $\frac{G_{100-positive} \cap G'_{100-positive}}{G'_{100-positive}}$ |
| Negative Precision@100 | $\frac{G_{100-negative} \cap G'_{100-negative}}{G'_{100-negative}}$ |

\* $n$ represents the number of landmark genes in expression profiles, $G$ represents the sets of top 100 positive/negative genes, $G'$ represents the sets of top 100 predicted genes.

In the context of chemical-blind splitting, different molecular representation and model initialization methods were tested. It was found that initializing the parameters of TranSiGen using perturbational profiles by gene knockdown leads to better performance compared to random initialization. Additionally, the pre-training representation using Knowledge-guided Pretraining of Graph Transformer (KPGT)^17^ further enhances the performance of inferring DEGs, surpassing the molecular fingerprint ECFP4 (Supplementary Table 1).

For cell-blind splitting, as the number of cells in the training set increases, TranSiGen demonstrates improved performance in inferring DEGs on seven unseen cells during training (Fig. 3d and Supplementary Table 3). However, its ability to generalize to new cells is not as evident as its ability to generalize to novel compounds that are not in the training set (Supplementary Table 1 and 3). This finding suggests that cross-cell prediction is challenging because both the cell type and its own status have a significant influence on the transcriptional profile. Effective prediction requires considering the basal profiles of the cells. Furthermore, this finding highlights the limitations in disregarding the impact of cell type on transcriptional profile prediction^10,18^.

In summary, TranSiGen stands out among other models in predicting DEGs in chemicalblind splitting and random splitting tests. It can also be utilized for inferring DEGs in cell-blind splitting. This further highlights the effectiveness of TranSiGen, which is based on self-supervised representation learning for transcriptional profiling.

### Ligand-based virtual screening with TranSiGen-derived representation

Compounds with shared mechanisms cause similar changes in cellular signaling, resulting in similar gene expression profiles^1,7,19^. As a simulation of chemical-induced transcriptional profiling, the TranSiGen-derived representation shows higher PCC among active compounds with the same target compared to the PCC between active and inactive compounds (Supplementary Fig. 4). In this section, the TranSiGen-derived representation was used to assess whether a compound is active against a specific target. Specifically, DEGs inferred by TranSiGen and other baseline models were used as the representations of compounds. Then, RF models based on these representations were constructed to screen active compounds for a target of interest.

Fig. 4a shows the performance of screening 5-hydroxytryptamine receptor 2A (HTR2A) active compounds on seven cell lines. The model based on TranSiGen-derived representation outperforms other perturbational representations by a significant margin. This result is further supported by the dimensionality reduction distribution of active/inactive compounds, where TranSiGen-derived representations clearly distinguish between the two, while other perturbational representations exhibit overlapped distributions (Fig. 4b and Supplementary Fig. 5).

**Fig. 4.**
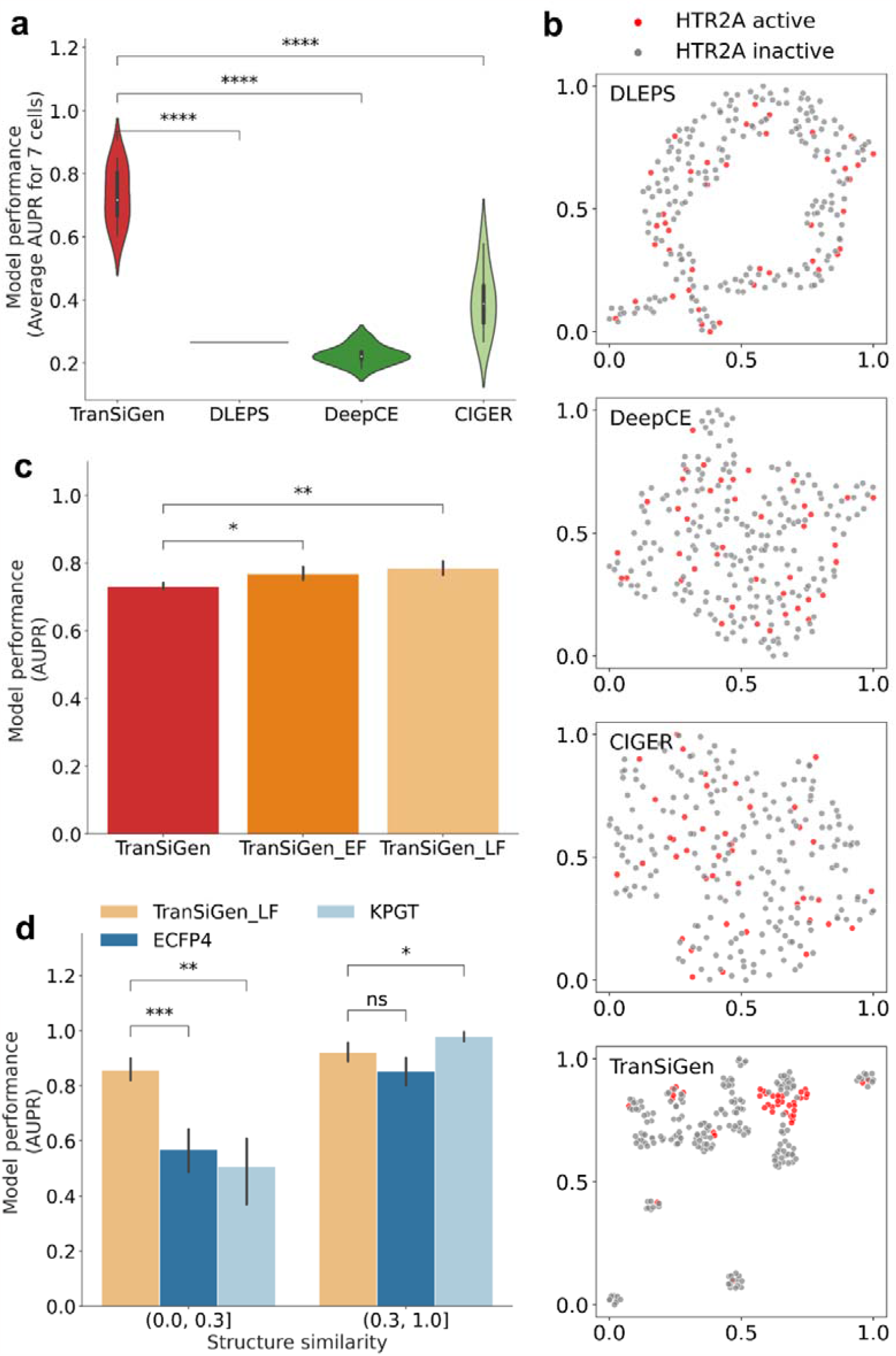
Model performance of ligand-based virtual screening on target HTR2A. **a** Performance of active compound prediction using different perturbational representations. **b** Dimensionality reduction visualization of HTR2A active and inactive compounds based on various inferred perturbational representations. **c** Performance of active compound prediction by applying early fusion and late fusion for TranSiGen-derived representation from seven different cell lines. **d** Performance of active compounds prediction within different thresholds of max similarity of test molecules relative to train data. Statistical t-test was applied between the models. (Note: ****, *p* < 0.0001; ***, 0.0001 < *p* ≤ 0.001; **, 0.001 < *p* ≤ 0.01; *, 0.01 < *p* ≤ 0.05 and ns, 0.05 < *p* ≤ 1.0)

TranSiGen can be used to obtain the characteristics of each compound in different cellular backgrounds. In this study, TranSiGen-derived representations from seven cell lines were fused to evaluate their impact on compound screening performance. Early fusion involves concatenating TranSiGen-derived representations from seven cells into one single feature, while late fusion merges the prediction results from seven cells. These two models are denoted as TranSiGen_EF and TranSiGen_LF for early and late fusion, respectively. It was observed that fusing TranSiGen-derived representations from different cell lines further enhances the screening performance of active compounds compared to individual cells alone (Fig. 4c). However, the performance improvement of TranSiGen_EF is not as significant as that of TranSiGen_LF, possibly due to the “curse of dimensionality”^20^. High-dimensional input features in TranSiGen_EF make it difficult to learn meaningful patterns. Similar phenomena are also observed in ligand-based virtual screening for other four targets evaluated (Supplementary Fig. 6).

As a molecular representation method, the TranSiGen-derived representation was compared to other molecular structural representations such as molecular fingerprint ECFP4 and the pre-trained representation KGPT. The maximum Tanimoto similarities of test molecules relative to training molecules were calculated using ECFP4. The performance of screening active compounds was evaluated at different maximal similarity thresholds. For compounds that are dissimilar to the training set (chemical structure similarity∈ (0.0, 0.3]), the TranSiGen-based model demonstrates better predictive ability than structure representation-based model (Fig. 4d and Supplementary Fig. 6). This suggests that using transcriptional profiling, such as TranSiGen-derived representation, may have advantages in screening for new scaffold compounds that differ from known compound structures.

Therefore, TranSiGen-derived representation can be used as a new form of molecular representation for describing the characteristics of compounds from various cell contexts. It can also complement the structure-based representation and offer advantages in ligand-based virtual screening.

### Drug response prediction with TranSiGen-derived representation

Chemical-induced transcriptional profiles directly associate molecular features with the cellular effect of a particular drug. This association is beneficial for characterizing drug response in different cells^21–23^. Here, we applied the TranSiGen-derived representation to predict the area under the dose-response curve (AUC) of a compound on a specific cell line. The AUCs were obtained from the cancer treatment response portal (CTRP)^24,25^. We defined compounds with AUCs ≥ 5.5 as resistant to cell lines, while those with AUCs < 5.5 were considered sensitive^24^. More details about the dataset can be found in Supplementary Table 4.

To determine whether compounds can be classified as sensitive or resistant to a specific cell line based on TranSiGen-derived representation, we first assessed the profiling similarity among different compounds. In Fig. 5a, we calculated the PCC within a group of sensitive compounds (denoted as Sensitive), as well as the PCC between sensitive and resistant compounds (denoted as Sensitive∼Resistant). Additionally, we compared the structural similarities of the two groups using Tanimoto similarity based on the molecular fingerprint ECFP4 (Fig. 5b). The results indicate that the Tanimoto similarities within Sensitive group and the Tanimoto similarities within Sensitive∼Resistant group are not significantly different on each cell line, suggesting that the structural representation ECFP4 cannot distinguish sensitive and resistant compounds (Fig. 5b). In contrast, we observed that the profiling similarities of Sensitive are significantly higher than those of Sensitive∼Resistant on most cell lines (Fig. 5a). This finding demonstrates the effective discrimination between sensitive and resistant compounds achieved through TranSiGen-derived representation.

**Fig. 5.**
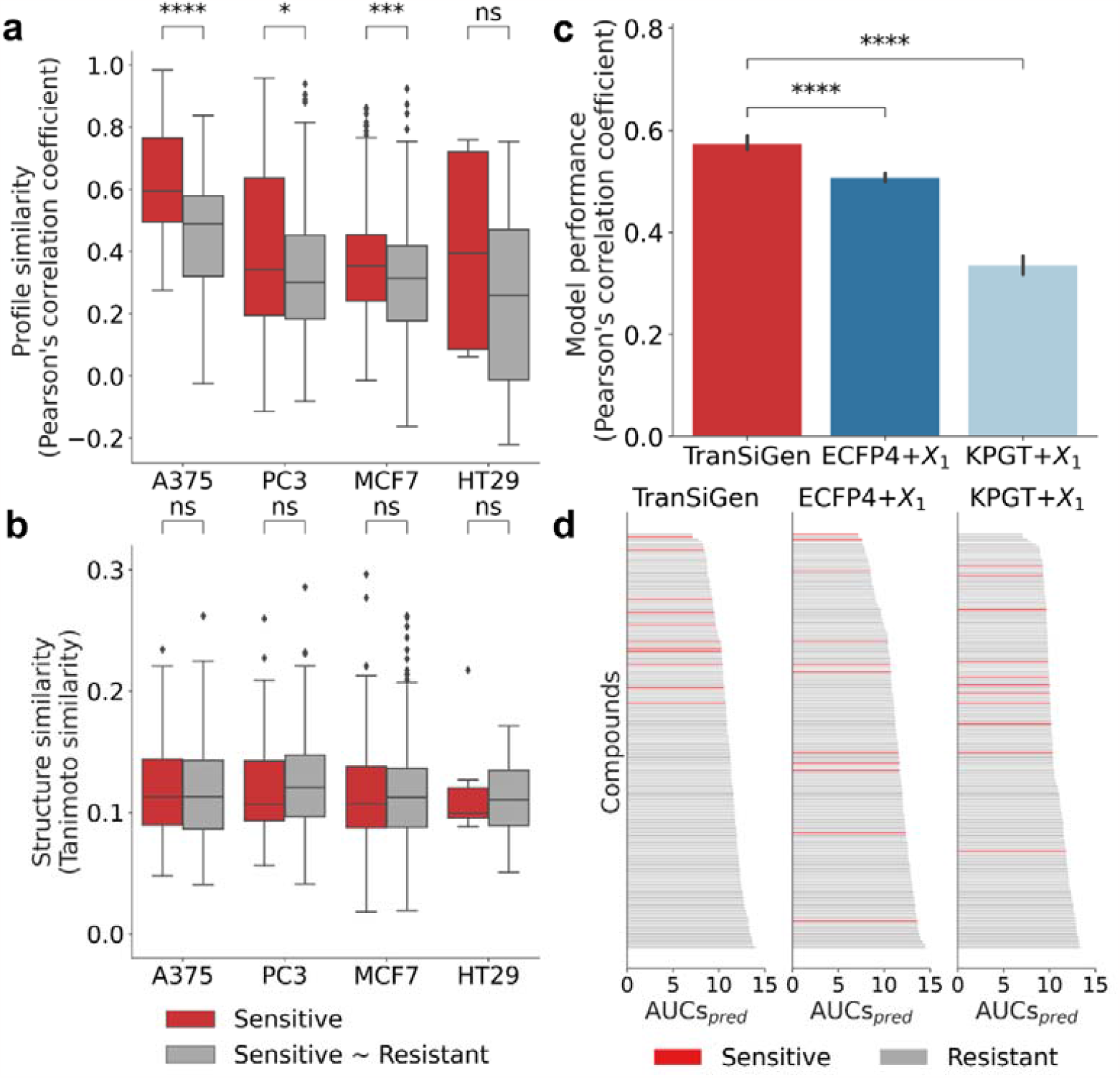
Model performance of drug response prediction. **a** The Pearson’s correlation coefficients within a group of sensitive compounds and the Pearson’s correlation coefficients between sensitive and resistant compounds based on TranSiGen-derived representation. **b** The Tanimoto similarity within a group of sensitive compounds and the similarity between sensitive and resistant compounds based on molecular fingerprint ECFP4, and the Mann-Whitney test was used to analyze the data. **c** Performance of predicting drug response using various type of representations, and statistical t-test was applied between the models. **d** Ranking results of compounds by AUCs_*pred*_ of models based on various type of representations. (Note: ****, *p* < 0.0001; ***, 0.0001 < *p* ≤ 0.001; **, 0.001 < *p* ≤ 0.01; *, 0.01 < *p* ≤ 0.05 and ns, 0.05 < *p* ≤ 1.0)

Furthermore, the TranSiGen-derived representation was used for drug response prediction in downstream task using a RF model. Its performance was compared with RF models based on other alternative representations, including perturbational representations generated by baseline models (DLEPS, DeepCE, and CIGER), as well as representations combining molecular structures and cell information (ECFP4+*X*_1_ and KPGT+*X*_1_). As shown in Fig. 5c and Supplementary Fig. 7a, the TranSiGen-based model demonstrates significantly better performance than other models. Additionally, to evaluate the screening performance, compounds were ranked by their predicted AUCs (AUCs_*pred*_), and classified as sensitive or resistant according to their true AUCs. The results showed that the TranSiGen-based model predicted sensitive compounds with smaller AUCs_*pred*_ and higher rankings, while other models ranked the sensitive compounds randomly (Fig. 5d and Supplementary Fig. 7b). This indicates that the TranSiGen-based model has superior screening ability for sensitive compounds.

In summary, the TranSiGen-derived representation, simulating DEGs of compounds on cell lines, exhibits a distinguishable feature for sensitive and resistant compounds and demonstrates outstanding performance on drug response prediction.

### Phenotype-based drug repurposing for the treatment of pancreatic cancer

Associating chemical-induced transcriptional profiles with diseases can help identify potential compounds for treating specific diseases^2,7^. TranSiGen-derived transcriptional profiles can be used alongside the profiles derived from chemical-treated and -untreated disease states to screen candidate compounds for disease treatment.

In this study, we integrated TranSiGen into a phenotype-based drug repurposing pipeline for pancreatic cancer^26^ to assess its ability to prioritize sensitive compounds for the YAPC pancreatic cancer cell line from a pool of 1,625 compounds in the PRISM Repurposing dataset^27^. We used two phenotype-based strategies and compared them to a conventional structural similarity-based protocol (Fig. 6a). TranSiGen_DRUG used the real DEGs of approved pancreatic cancer drugs to identify compounds with similar perturbation effects. Conversely, TranSiGen_DISEASE looked for compounds that can reverse the DEGs of pancreatic cancer. Both strategies used connectivity scores^28^ to measure the relationship between DEGs. For comparison, ECFP4_DRUG was implemented to find compounds structurally similar to the approved drugs using ECFP4-based Tanimoto similarity. For more information, please refer to the section “Phenotype-based drug repurposing for pancreatic cancer” in the Methods section.

**Fig. 6.**
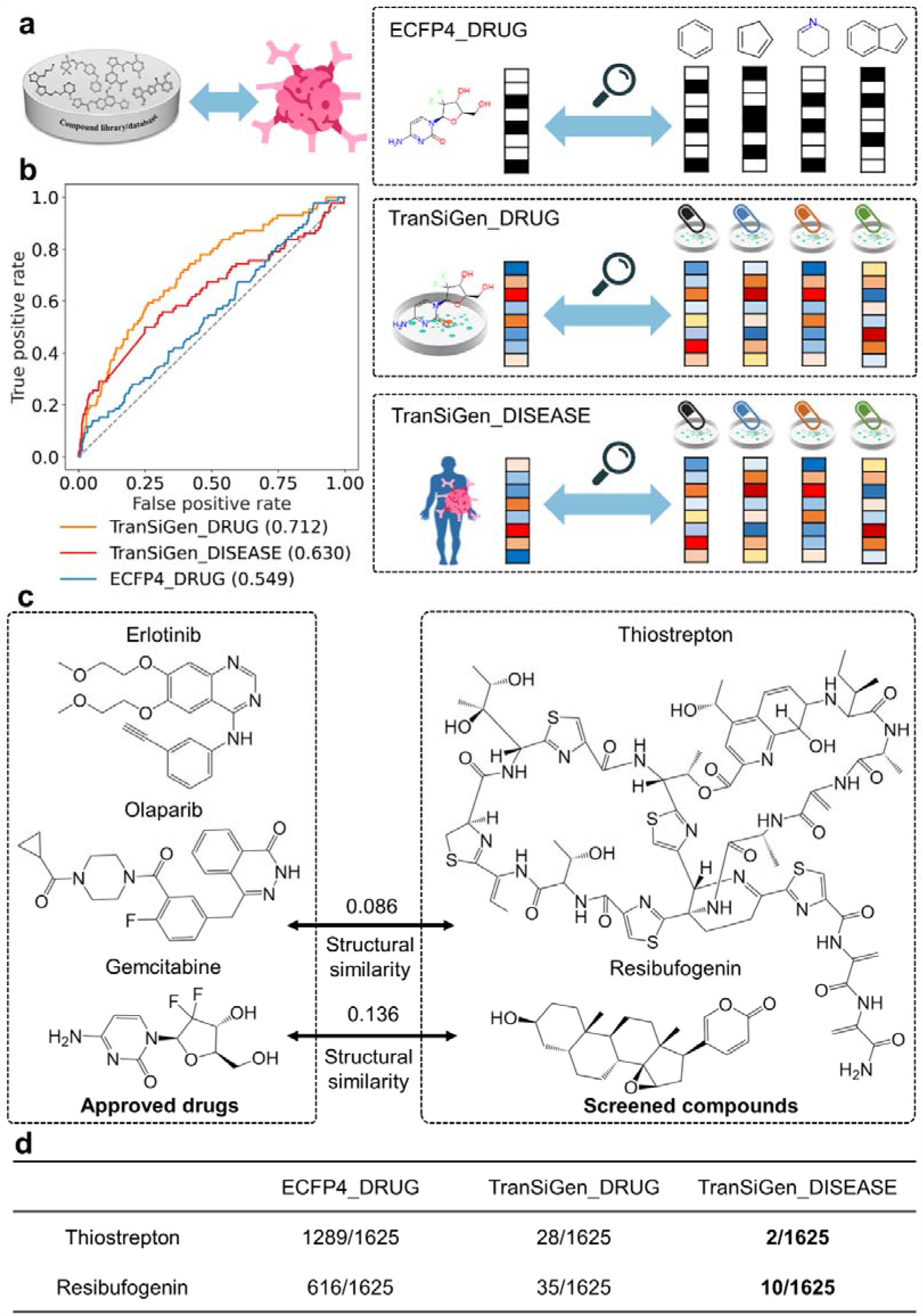
Phenotype-based drug repurposing for the treatment of pancreatic cancer. **a** The flow chart of drug repurposing strategy. **b** The screening performance of phenotype-based strategy and structural similarity-based strategy. **c** TranSiGen_DISEASE screened compounds that are capable of inhibiting pancreatic cancer cells, and their max structural similarities to approved drugs. **d** The rankings of thiostrepton and resibufogenin from different screening strategies.

The screening performance of the three methods is shown in Fig. 6b. ECFP4_DRUG yielded the worst predictive classification performance, followed by TranSiGen_DISEASE, and the best was TranSiGen_Drug. Notably, the TranSiGen_DISEASE approach doesn’t require any chemical-treated profiles, simulating scenarios where diseases lack known therapeutic drugs. This is a challenge not addressed by the structural similarity-based strategy. Even without using perturbed profiles of known drugs, TranSiGen_DISEASE effectively enriched hits among the top-ranking compounds (Supplementary Table 5). Phenotype-based strategies can identify compounds less similar to the approved drugs than those screened by ECFP4_DRUG (Fig. 6c and Supplementary Fig. 8). For instance, nature products thiostrepton and resibufogenin (Fig. 6c) ranked in the top 10 without sensitive annotations in PRISM dataset (Supplementary Table 6). Their abilities to inhibit pancreatic cancer cells have been confirmed by a literature survey^29,30^. Thiostrepton, a natural cyclic oligopeptide, reduces the viability and clonogenicity of pancreatic cancer cell lines and induces ferroptosis via STAT3/GPX4 signalling^30^. Resibufogenin, a steroid lactone from the skin venom gland of toads, demonstrated potent anti-pancreatic cancer effects in vivo and in vitro, and can induce caspase-dependent apoptosis^29^. Fig. 6d summarizes the rankings of these two compounds in different screening strategies. Both phenotype-based strategies, TranSiGen_DISEASE and TranSiGen_DRUG, consistently prioritized them. In contrast, the structure-based strategy ECFP4_DRUG failed to effectively prioritize these nature products, ranking them at 1,289 and 616, respectively. This may be attributed to the large structural differences between them and approved drugs, highlighting the inherent limitation of a structure-based strategy.

These results highlight the effectiveness of the phenotype-based strategies that use TranSiGen-derived representation in identifying potent candidate compounds, including those with unique structures. Overall, TranSiGen expands the range of compounds that can be screened with predicted transcriptional perturbational profiles. It can be easily integrated into a phenotype-based drug repurposing pipeline, improving drug discovery efficiency and minimizing costs.

## Discussion

Given the widespread application of perturbational gene expression profiling in biomedical research and the limitations arising from random noise and systemic biases in profiling data, we propose TranSiGen, a VAE-based model. TranSiGen employs a self-supervised representation learning strategy to overcome these inherent limitations of data and infer novel chemical-induced gene expression profiles. TranSiGen excels in denoising and reconstructing the transcriptional profiling, as well as effectively learning cellular and compound features from data. It outperforms existing models in both random splitting and chemical-blind splitting, and also shows potential for inferring DEGs in new cell lines, a capability absents in existing models. Moreover, TranSiGen-derived profiles have served as a unified and standardized representation of phenotypic information, demonstrating its effectiveness in various downstream tasks, including ligand-based virtual screening, drug response prediction, and phenotype-based drug repurposing.

Therefore, TranSiGen efficiently learns the representation of transcriptional data and can predict novel chemical-induced transcriptional changes in a given cell line. We believe that integrating it into the pipeline of drug discovery and mechanism of action research can enhance the efficiency and reduce the costs, thereby promoting the development of biomedicine.

## Methods

### Transcriptional data processing

LINCS^2^ has made publicly available resources on high-throughput gene expression profiles of different perturbagens, such as small molecule compounds and shRNAs. Using the L1000 assay, it is possible to measure the expression values of only 978 landmark genes, and while still recovering most of the full transcriptome. The latest CMAP LINCS 2020 dataset was used in this study^15^.

In LINCS, there are 5 levels of data. For this study, we utilized level 3 data, which included both raw perturbation profiles and control profiles (denoted as *X*_1_ and *X*_2_, respectively). To filter the profiles, we used the most common condition with a duration of 24 h and a dosage concentration of 10 μM for perturbed expression profiles by compounds. Additionally, we matched the expression profile with DMSO vehicle to the perturbed profiles on the same plate to create paired profiles *X*_1_ ∼ *X*_2_, minimizing batch effects between cases and controls. The extracted dataset contained 219,650 *X*_1_ ∼ *X*_2_ pairs for 8,316 compounds on 164 cell lines. Furthermore, MODZ was applied to ensure that only one *X*_1_ ∼ *X*_2_ pair per compound was included for each cell line with multiple *X*_1_ ∼ *X*_2_ pairs. The processed dataset contained the transcriptional profiles of 8,316 compounds on 164 cell lines, including 78,569 *X*_1_ ∼ *X*_2_ pairs.

The gene expression profiles induced by shRNA were processed using the same method described above. Profiles from the 10 most common cell lines (A375, A549, ASC, HA1E, HCC515, HT29, MCF7, NPC, PC3, and VCAP) measured after 24 h were selected. The control profile with an empty vector in the same plate was then paired with the perturbed profiles. The final dataset contained 188,509 *X*_1_ ∼ *X*_2_ pairs consisting of 4,112 shRNAs on 10 cell lines, which were used to initialize two VAEs in TranSiGen.

### Compound representations

Considering that the current number of compounds with experimentally measured gene expression profiles is still limited compared to the vast chemical space, TranSiGen utilized the pre-trained molecular representation KPGT^17^ for compounds. KPGT is a novel self-supervised learning framework for molecular graph representation. It leveraged a knowledge-guided pre-training strategy to capture rich structural and semantic information from large-scale unlabeled molecular graphs. In this study, the 2034-dimensional representation obtained from the KPGT pre-trained model was used as the molecular input for TranSiGen.

Alternatively, chemical fingerprints, widely used as a form of molecular representation in machine learning, was also used as molecular input for TranSiGen. There are represented as binary vectors indicating the presence or absence of particular substructures in compounds. Specifically, the molecular fingerprint ECFP4^31^ with a radius of 2 and a length of 2048 was used here.

### TranSiGen architecture

The VAE^32^ is a deep generative model consisting of an encoder and a decoder. VAE is capable of learning an efficient and meaningful latent space from high-dimensional data by compressing and reconstructing the original input. Unlike the standard autoencoder, which maps the input to a point in the latent space and trains by minimizing the reconstruction error, VAE encodes the input to a distribution. This requires the addition of a Kullback-Leibler (KL) divergence term to the reconstruction loss, which constrains the latent vectors to match a Gaussian distribution.

The architecture of TranSiGen consists of two VAEs: one for encoding the basal profiles *X*_1_ and the other for encoding the perturbation profiles *X*_2_. TranSiGen minimizes the loss of learning the representations of *X*_1_ and *X*_2_. Additionally, a linear function is used to map from the latent representation of and the hidden representation of the compound representation *C*_*mol*_ to the perturbed latent representation *Z*_2_*FZ*_1_ of 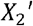, mimicking the chemical-induced transcription changes. The layer and dimension details of TranSiGen are shown in Supplementary differential expression genes Δ*X*^′^. This involves minimizing the reconstruction loss between Fig. 1. During the training process, TranSiGen also minimizes the loss of predicting the *X*_2_ ^′^ − *X*_1_ and *X*_2_ − *X*_1_, as well as constraining the predicted perturbed latent representation *Z*_2_*FZ*_1_ match to the latent representation *Z*_2_. The loss function of TranSiGen is defined as follow:

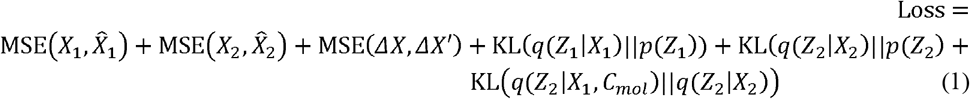

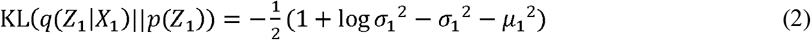

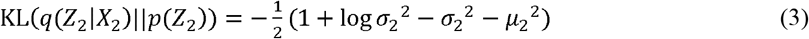

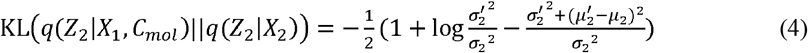

where *μ*_1_ and 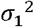 represent the mean and variance for *q*(*Z*_1_|*X*_1_), μ_2_ 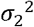 and represent the mean and variance for *q*(*Z*_1_|*X*_1_), *μ*_2_, 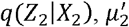 and 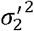 represent the mean and variance for *q*(*Z*_2_|*X*_1_, *C*_*mol*_) .

### Performance evaluation metrics

As shown in Table 2, the model’s prediction performance was evaluated using following metrics: Root mean squared error (RMSE), Pearson’s correlation coefficient, and Precision@K. RMSE and Pearson coefficient were used to measure the prediction performance on the overall landmark genes. Precision@k, on the other hand, focused on the most significantly up- and down-regulated expressed genes. In this study, Positive Precision@100 was evaluated for the top 100 up-regulated genes, while Negative Precision@100 was evaluated for the top 100 down-regulated genes.

### Ligand-based virtual screening

For compounds with transcriptional profiles, the target annotations were collected from the compound information file provided by LINCS2020^15^ and PubChem^33^ (https://pubchem.ncbi.nlm.nih.gov). Specifically, the targets of the compound were determined according to IC_50_, K_i_, K_d_ less than 10 μM in PubChem, and the target information from these two sources was deduplicated and integrated.

In the task of ligand-based virtual screening, compounds with target annotations were selected from 7 cell lines (A375, HELA, PC3, MCF7, HT29, YAPC, and HA1E). Among these, the target HTR2A had the most active compounds (Supplementary Fig. 3), and was selected for the task. Inactive compounds were sampled from the remaining compounds at a ratio of 1:5 to construct a dataset for the active compound screening. The compounds from each cell line were randomly split into training and test set with a ratio of 4:1.

Given the limited availability of dataset for active compound screening, we used RF for active compound prediction. To construct the RF classifiers, we used two types of features: inferred perturbational representations (TranSiGen, DLEPS, DeepCE, and CIGER) and structural representations (molecular fingerprint ECFP4, and pretrained representation KPGT). We conducted a hyperparameter search for “n_estimators”, “max_depth”, “criterion” and “obb_score” to find the optimal model. To evaluate the model performance, we used the area under the Precision–Recall curve (AUPR). The training-evaluation procedure was repeated five times with different random seeds to determine the model performance. These processes were implemented using scikit-learn^34^.

### Drug response prediction

The CTRP^24,25^ is a widely used cancer cell response dataset that associate genetic, lineage, and other cellular molecular characteristics of cancer cell lines with drug sensitivity. It quantitatively profiles the sensitivity of cancer cell lines to small molecules. The AUC label is a dose-independent measure of compound sensitivity. Smaller AUCs indicate greater sensitivity of cells to the drugs. A subset of the drug response dataset for 267 compounds on four cell lines (A375, PC3, MCF7, and HT29) was obtained from CTRP, and the details of the dataset are shown in Supplementary Table 4. The processed dataset was split into training and test set at 4:1 ratio by compounds.

Similarly, RF regression models were used to predict drug response. Inferred perturbational representations (TranSiGen, DLEPS, DeepCE, and CIGER) and representations combining molecular structures and cell information (ECFP4+*X*_1_ and KPGT+*X*_1_) were used. Four hyperparameters, including “n_estimators”, “max_depth”, “criterion” and “obb_score”, were considered to obtain the optimal model. The model’s performance was evaluated by the Pearson’s correlation coefficient. The model’s performance was assessed by repeating training-evaluation procedure five times with different random seeds.

### Phenotype-based drug repurposing for pancreatic cancer

#### Differential gene expression profiles of approved drugs

The approved drugs for pancreatic cancer were downloaded from https://www.cancer.gov/about-cancer/treatment/drugs/pancreatic. Among them, the DEGs of erlotinib, olaparib and gemcitabine were obtained from LINCS 2020 dataset^15^. These profiles were used for subsequent phenotype-based drug repurposing for pancreatic cancer.

#### Differential gene expression profile of disease

The pancreatic adenocarcinoma cohort of the The Cancer Genome Atlas (TCGA)^35^ was downloaded from UCSC Xena (https://xenabrowser.net/). This cohort includes RNA-seq expression data of tumor samples and normal samples. The DESeq2^36^ method was used to analyze the differential gene expression for pancreatic cancer. DEGs for pancreatic cancer were selected based on the following criteria: |log2Foldchange|>□1.5, p-□ <□0.05 and false discovery rate< 0.25. A total of 293 up-regulated genes and 168 down-regulated genes were identified.

#### Inferring perturbation gene expression profiles of compounds

This study utilized the PRISM Repurposing dataset^27^, which includes primary and secondary screening datasets, for phenotype-based repurposing for pancreatic cancer. The compounds from PRISM secondary screen were evaluated based on their AUC values, which indicate compound sensitivities on cells and serve as labels for screening performance assessment.

The dataset was downloaded from https://depmap.org/repurposing/. Compounds with ground-truth expression profiles in the LINCS 2020 dataset were excluded, resulting a dataset contains 1,625 compounds. TranSiGen inferred the DEGs Δ *X* ′ of 978 landmark genes associated with these compounds in YAPC pancreatic cancer cell. Additionally, and the expression values of 9,196 best inferred genes were inferred from the generated 978 landmark genes to obtain the predicted expression values of 10,174 genes. The inference weight matrix was obtained from the L1000 project^2^.

### Connectivity score

The connectivity score, obtained from the gene set enrichment analysis^28^, is used to measure the relationship between transcriptional profiles. The connectivity score ranges from -1 to 1, where -1 indicates a complete reversal of the query profile to the reference profile, while 1 indicates a complete similarity of the query and the reference profile.

Firstly, the enrichment score (ES) is used to evaluate the enrichment of a predefined gene set at the top or bottom of the reference differential gene list. The enrichment scores for up-regulated gene set and down-regulated gene set are denoted as *a* and *b*, respectively:

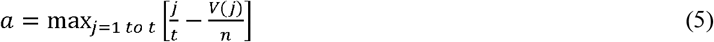

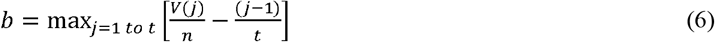

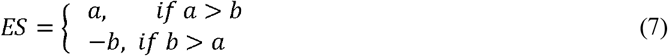

where *n* represents the number of genes in the expression profiles, *t* represents the number of genes in the predefined gene set, and *V*(*j*)represents the rank of a specific gene in the rank list.

Next, the above equations are used to calculate the enrichment scores of the predefined up-regulated and down-regulated genes by the query profile, resulting in the values *ES*_*up*_ and *ES*_*down*_. Finally, considering these two enrichment scores together, the connectivity score of the query profile relative to the reference profile is calculated as follows:

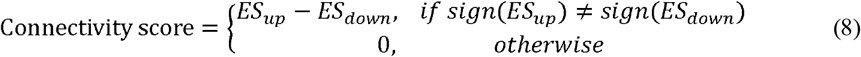

Specifically, the reference profiles consist of the inferred DEGs of all compounds from TranSiGen. The DEGs of approved drugs for pancreatic cancer were used to identify compounds with positive connectivity scores, while the DEGs of pancreatic cancer were employed to identify compounds with negative connectivity scores. In the screening dataset, the top 20% compounds having the lowest AUCs on the YAPC pancreatic cancer cell line were identified as hits, and the area under the ROC curve was used to evaluate the screening performance.

## Data availability

The expanded CMap LINCS Resource 2020 is available at https://clue.io/data/CMap2020#LINCS2020. The PRISM Repurposing dataset is available at https://depmap.org/repurposing/, and the pancreatic adenocarcinoma cohort of the TCGA is available https://xenabrowser.net/.

## Code availability

The code for model training and analysis is available at: https://github.com/myzheng-SIMM/TranSiGen, and have been deposited fully in the Zenodo under accession code https://zenodo.org/records/10056517.

## Acknowledgements

We gratefully acknowledge financial support from National Natural Science Foundation of China (T2225002, 82273855 to M.Y.Z. and 82204278 to X.T.L), SIMM-SHUTCM Traditional Chinese Medicine Innovation Joint Research Program (E2G805H), Shanghai Municipal Science and Technology Major Project, National Key Research, and Development Program of China (2022YFC3400504 to M.Y.Z.), the Youth Innovation Promotion Association CAS (2023296 to S.L.Z).

## Contributions

M.Z. and X.L. conceived the project and were responsible for the decision to submit the manuscript; X.T. implemented the TranSiGen model, conducted computational analysis and wrote the paper; N.Q, X.K., S.N., K.W., L.Z, Y.W. and S.Z. discussed the results and commented on the manuscript.

## Ethics declarations

