## Supplementary for "TranSiGen: Deep representation learning of chemical-induced transcriptional profile"

Supplementary Figures

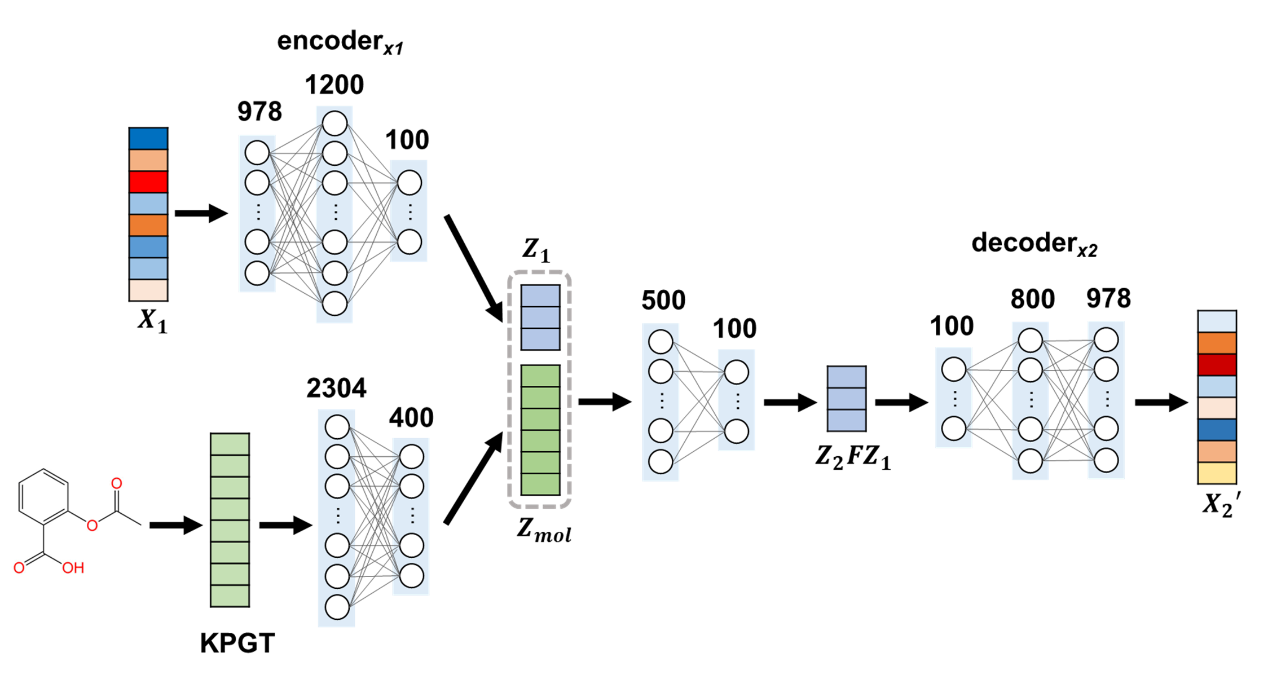

**Supplementary Fig. 1** Details of TranSiGen.

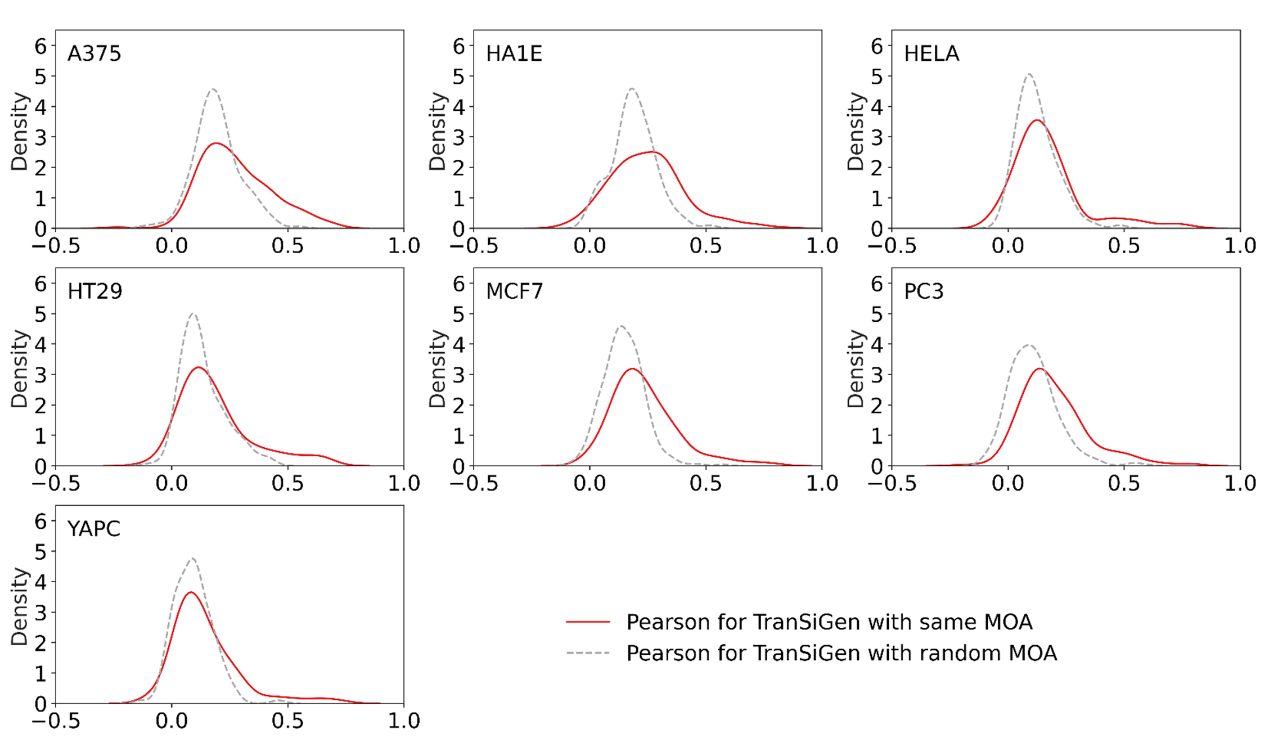

**Supplementary Fig. 2** Distribution of Pearson’s correlation coefficients of profiles for the same and random mechanism of action by TranSiGen-derived representation.

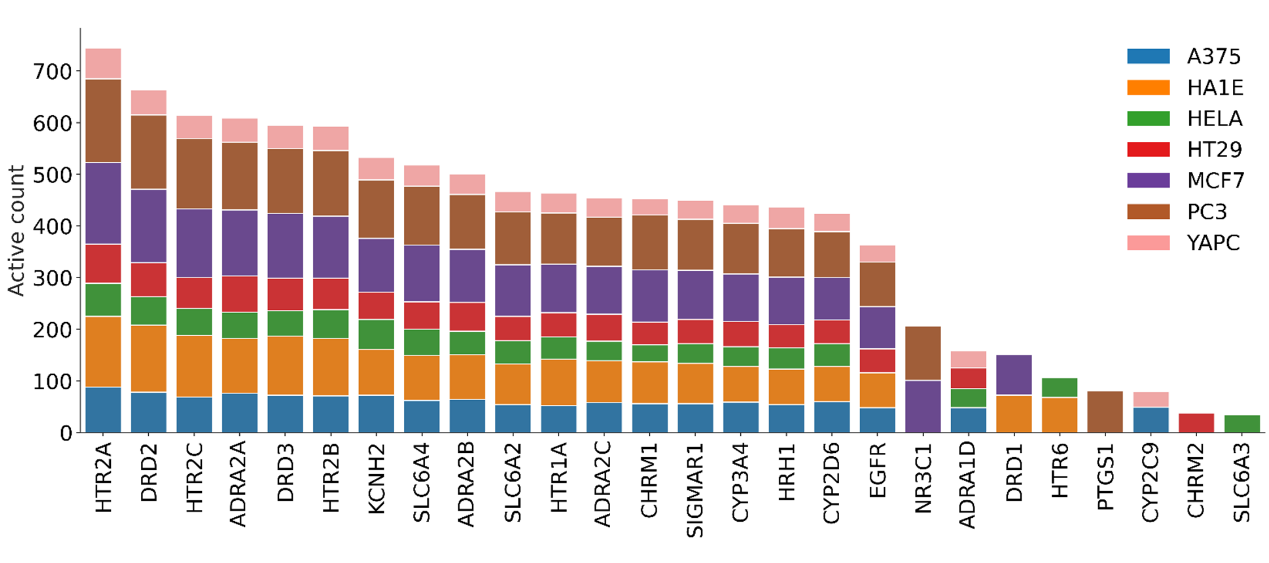
 **Supplementary Fig. 3** The number of active compounds on each target from different cell lines.

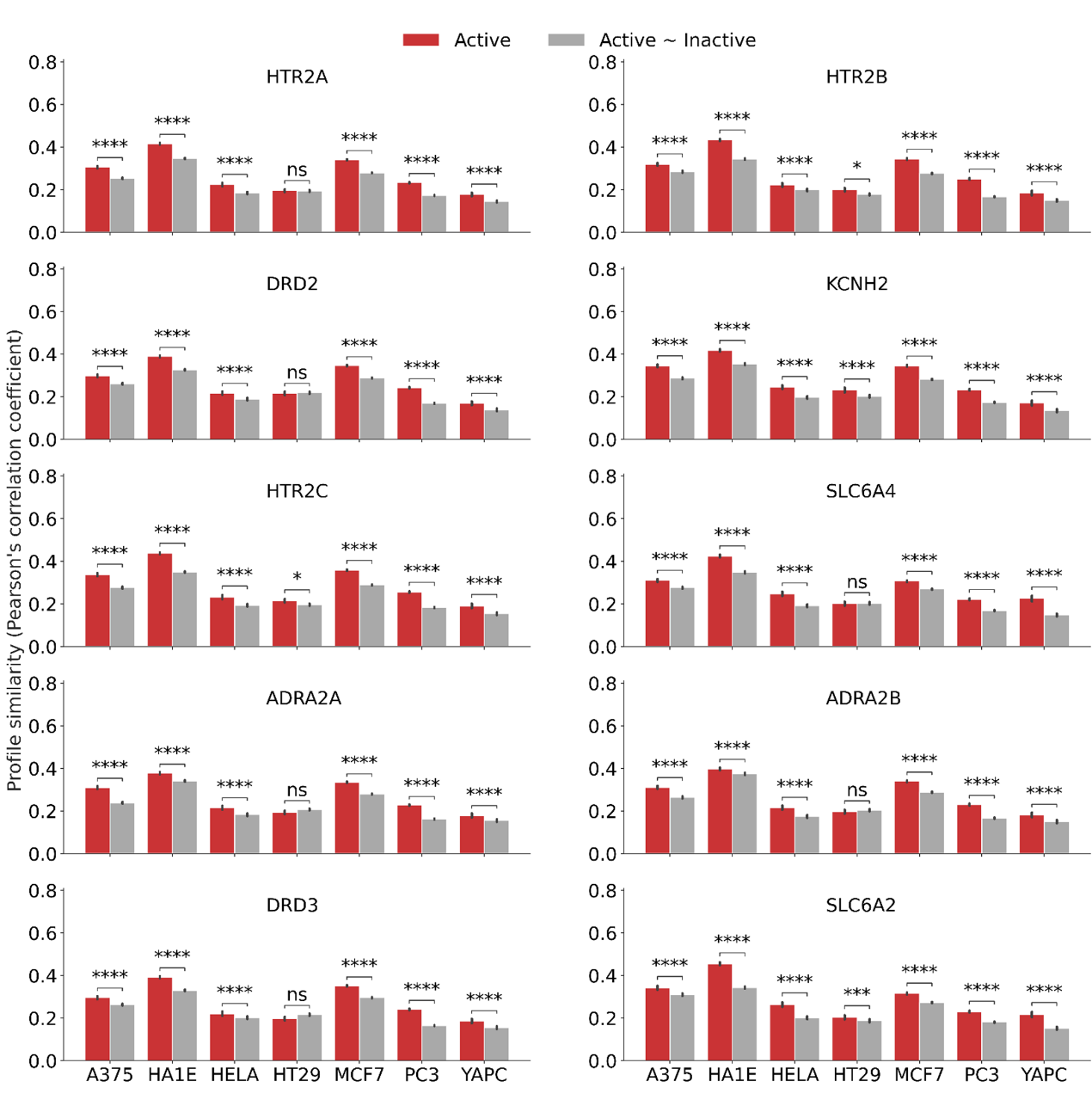

**Supplementary Fig. 4** The Pearson’s correlation coefficients within a group of active compounds and the Pearson’s correlation coefficients between active and inactive compounds based on TranSiGen-derived representation. The Mann-Whitney test was used to analyse the data. (Note: ∗∗∗∗, *p* < 0.0001; ∗∗∗, 0.0001 < *p* ≤ 0.001; ∗∗, 0.001 < *p* ≤ 0.01; ∗, 0.01 < *p* ≤ 0.05 and ns, 0.05 < *p* ≤ 1.0)

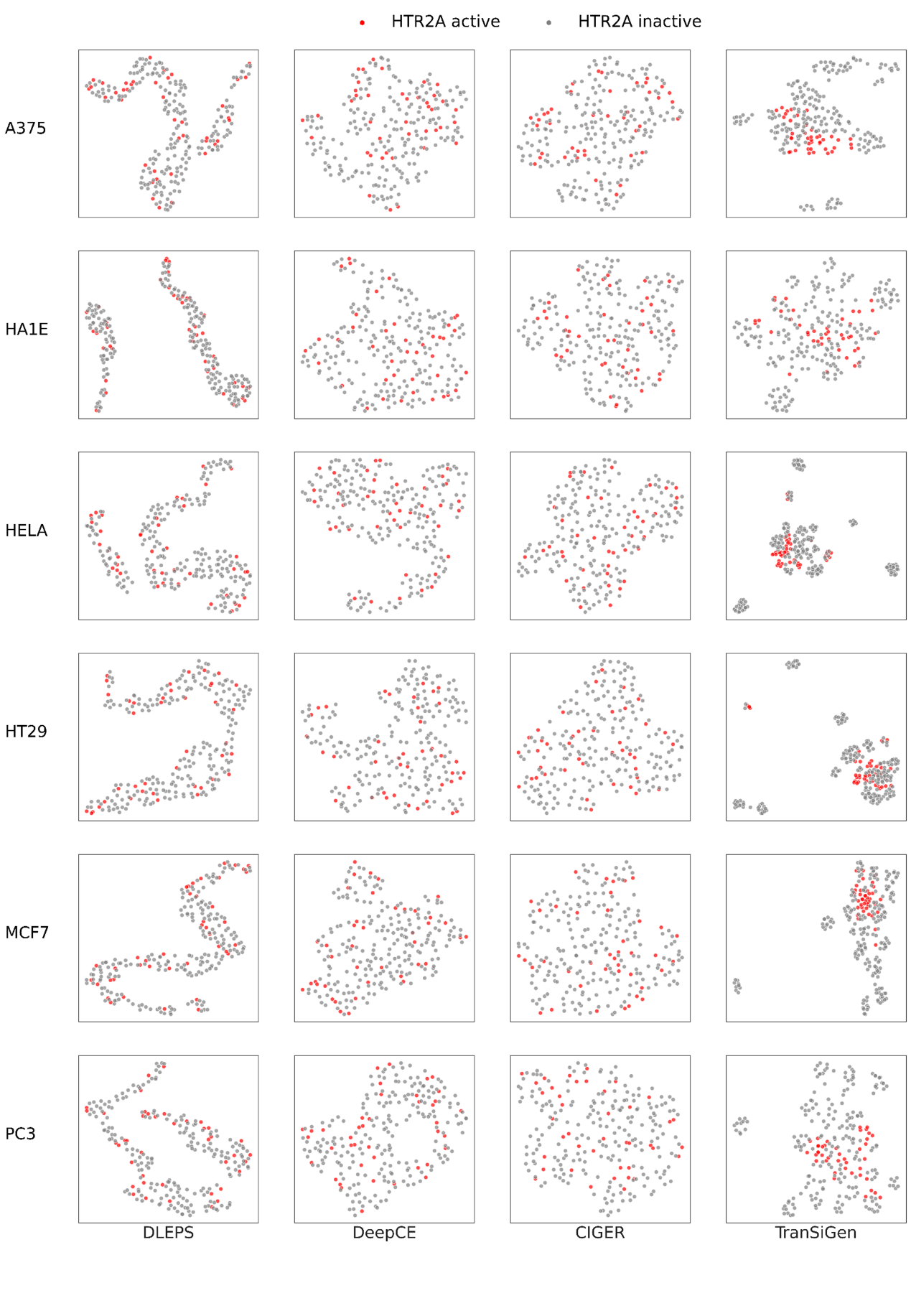

**Supplementary Fig. 5** Dimensionality reduction visualization of HTR2A active and inactive compounds based on various inferred perturbational representations.

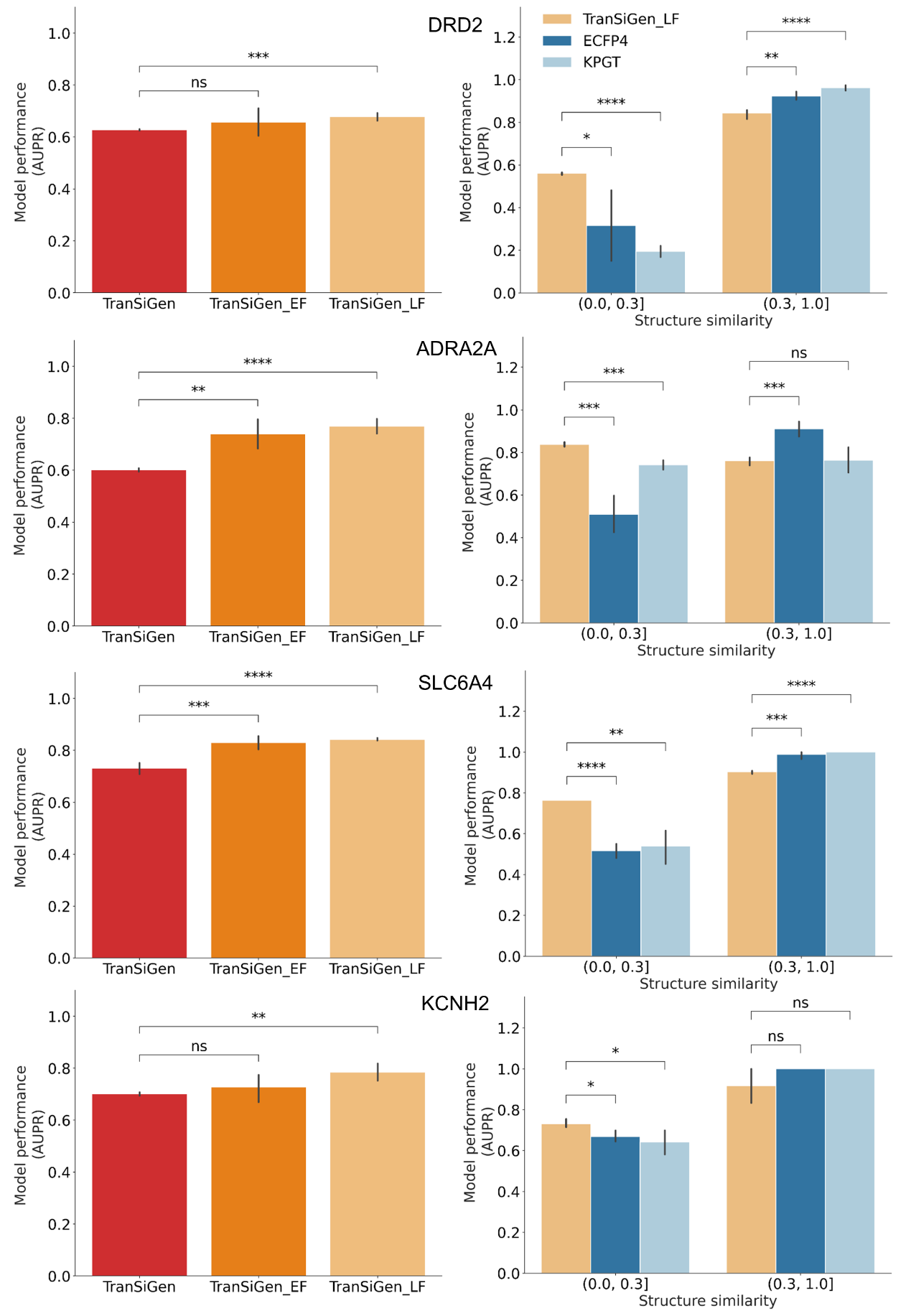

**Supplementary Fig. 6** Model performance of ligand-based virtual screening on target DRD2, ADRA2A, SLC6A4 and KCNH2. Statistical t-test was applied between the models. (Note: ∗∗∗∗, *p* < 0.0001; ∗∗∗, 0.0001 < *p* ≤ 0.001; ∗∗, 0.001 < *p* ≤ 0.01; ∗, 0.01 < *p* ≤ 0.05 and ns, 0.05 < *p* ≤ 1.0)

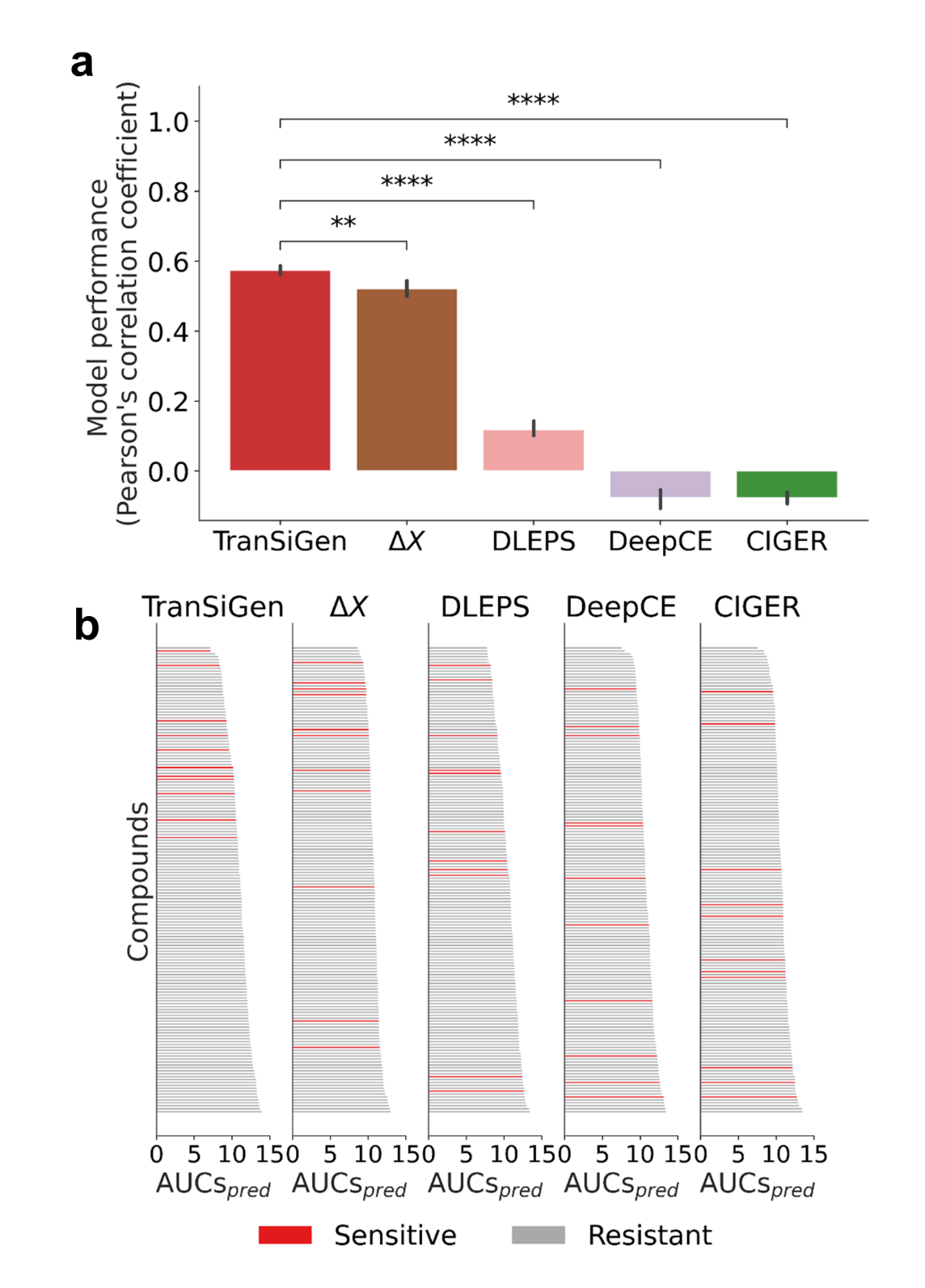

**Supplementary Fig. 7** Model performance of drug response prediction. a Performance of predicting drug response using various type of representations. b Ranking results of compounds by AUCs*_pred_* of models based on various type of representations. Statistical t-tests were applied between the models. (Note: ∗∗∗∗, *p* < 0.0001; ∗∗∗, 0.0001 < *p* ≤ 0.001; ∗∗, 0.001 < *p* ≤ 0.01; ∗, 0.01 < *p* ≤ 0.05 and ns, 0.05 < *p* ≤ 1.0)

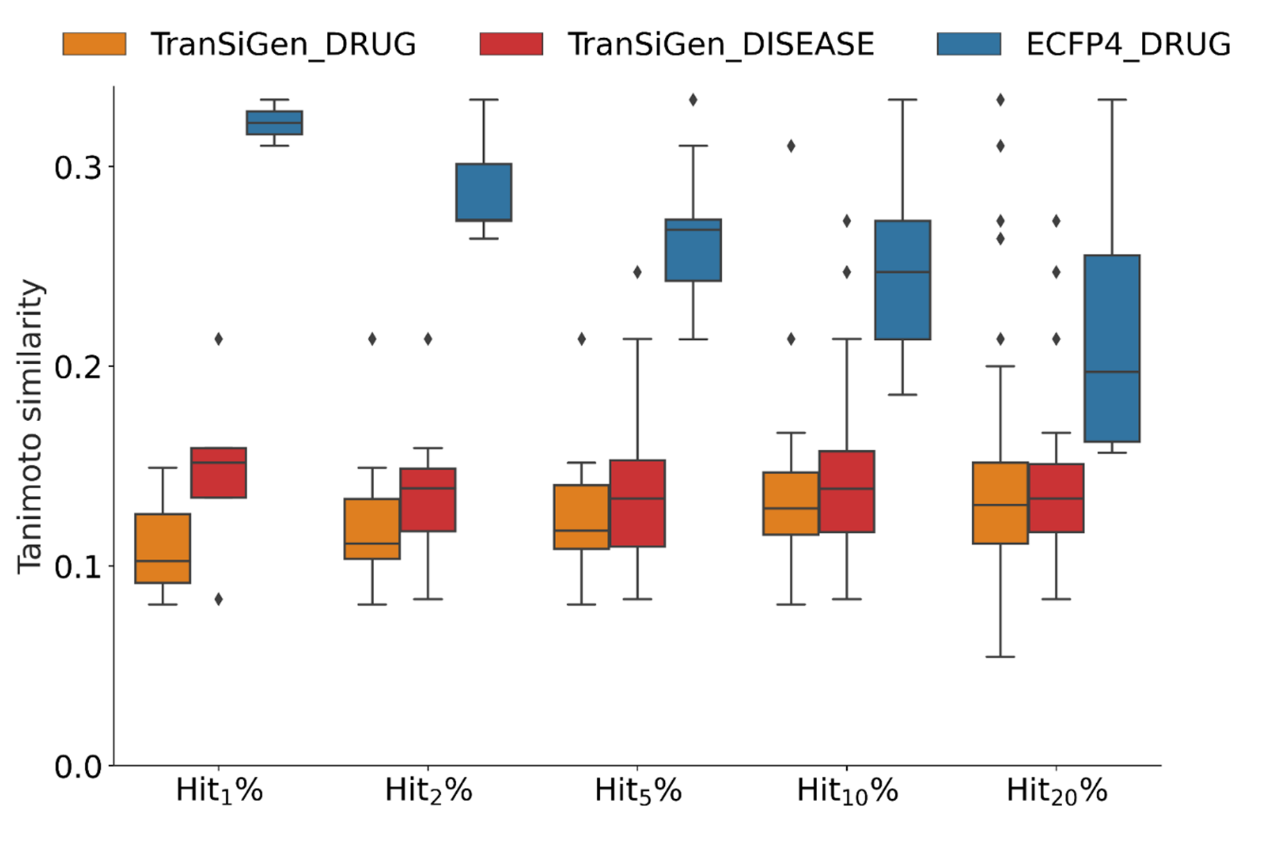

**Supplementary Fig. 8** The max similarity of hit compounds screened by phenotype-based strategy and structural similarity-based strategy to the approved drugs.

Supplementary Tables

**Supplementary Table 1.** Model performance for inferring DEGs in chemical-blind setting.

| Model | RMSE | Pearson | Positive P@100 | | Negative P@100 |
| --- | --- | --- | --- | --- | --- |
| DLEPS | 1.551±0.005 | 0.418±0.004 | | 0.244±0.002 | 0.316±0.002 |
| DeepCE | 1.773±0.034 | 0.430±0.020 | | 0.238±0.003 | 0.290±0.010 |
| CIGER | 2.551±0.282 | 0.436±0.002 | | 0.258±0.017 | 0.287±0.001 |
| TranSiGen  (ECFP4) | 0.661±0.003 | 0.517±0.001 | | 0.363±0.002 | 0.387±0.003 |
| TranSiGen  (KPGT) | 0.641±0.002 | 0.540±0.002 | | 0.381±0.002 | 0.397±0.001 |
| TranSiGen  (ECFP4; full data; init_random) | 0.527±0.004 | 0.609±0.001 | | 0.433±0.002 | 0.442±0.000 |
| TranSiGen  (ECFP4; full data) | 0.522±0.005 | 0.615±0.002 | | 0.441±0.000 | 0.448±0.004 |
| TranSiGen  (KPGT; full data;  init_random) | 0.524±0.004 | 0.613±0.001 | | 0.437±0.002 | 0.445±0.002 |
| TranSiGen  (KPGT; full data) | **0.520±0.003** | **0.619**±**0.000** | | **0.443±0.003** | **0.452±0.002** |

**Supplementary Table 2.** Model performance for inferring DEGs in random splitting.

| Model | RMSE | Pearson | Positive P@100 | Negative P@100 |
| --- | --- | --- | --- | --- |
| DeepCE | 1.648±0.002 | 0.519±0.002 | 0.286±0.002 | 0.329±0.002 |
| CIGER | 3.421±1.248 | 0.512±0.006 | 0.300±0.008 | 0.326±0.007 |
| TranSiGen (ECFP4) | 0.615±0.008 | 0.626±0.005 | 0.449±0.005 | 0.465±0.005 |
| TranSiGen (KPGT) | 0.610±0.006 | 0.631±0.006 | 0.453±0.008 | 0.467±0.005 |
| TranSiGen (ECFP4; full data) | 0.502±0.001 | 0.639±0.001 | **0.460±0.001** | 0.469±0.001 |
| TranSiGen (KPGT; full data) | **0.501±0.002** | **0.640±0.001** | 0.459±0.001 | **0.471±0.001** |

**Supplementary Table 3.** Model performance for inferring DEGs in cell-blind splitting.

| Model | RMSE | Pearson | Positive P@100 | Negative P@100 |
| --- | --- | --- | --- | --- |
| TranSiGen (KPGT;10 cells) | 1.198±0.004 | 0.260±0.001 | 0.214±0.001 | 0.200±0.002 |
| TranSiGen (KPGT;50 cells) | 1.073±0.008 | 0.297±0.003 | 0.241±0.002 | 0.223±0.004 |
| TranSiGen (KPGT;150 cells) | **0.960±0.012** | **0.324**±**0.003** | **0.256**±**0.002** | **0.232**±**0.004** |

**Supplementary Table 4.** Details of drug response dataset collected from CTRP.

| Cell | Compound | AUC | AUC<5.5 | AUC≥5.5 |
| --- | --- | --- | --- | --- |
| PC3 | 212 | 212 | 12 | 200 |
| MCF7 | 208 | 208 | 21 | 184 |
| A375 | 204 | 204 | 14 | 190 |
| HT29 | 179 | 179 | 4 | 175 |

**Supplementary Table 5.** The screening performance of phenotype-based strategy and structural similarity-based strategy.

|  | EF_1%_ | EF_2%_ | EF_5%_ | EF_10%_ | EF_20%_ |
| --- | --- | --- | --- | --- | --- |
| TranSiGen_DISEASE | **5.905** | **6.495** | **4.666** | **2.916** | 1.977 |
| TranSiGen_DRUG | 3.543 | 4.133 | 3.732 | 2.799 | **2.384** |
| ECFP4_DRUG | 2.362 | 3.543 | 2.333 | 1.516 | 1.337 |

**Supplementary Table 6.** Details of the top 20 candidate compounds by TranSiGen_DISEASE screening.

| Rank | Name | Connectivity score | AUC in PRISM | Target | MOA |
| --- | --- | --- | --- | --- | --- |
| 1 | SB-939 | -0.333 | 0.660 | HDAC1, HDAC3, HDAC4, HDAC5, HDAC9, HDAC10 | HDAC inhibitor |
| 2 | thiostrepton | -0.325 |  | FOXM1 | FOXM1 inhibitor, protein synthesis inhibitor |
| 3 | panobinostat | -0.323 | 0.728 | HDAC1, HDAC2, HDAC3, HDAC4, HDAC6, HDAC7, HDAC8, HDAC9 | HDAC inhibitor |
| 4 | belinostat | -0.319 | 0.874 | HDAC1, HDAC2, HDAC3, HDAC4, HDAC5, HDAC6, HDAC7, HDAC8, HDAC9, HDAC10, HDAC11, | HDAC inhibitor |
| 5 | dacinostat | -0.288 | 0.834 | HDAC1, HDAC2, HDAC3, HDAC4, HDAC5, HDAC6, HDAC7, HDAC8, HDAC9 | HDAC inhibitor |
| 6 | BNTX | -0.287 | 0.823 | OPRD1, OPRK1, OPRM1 | opioid receptor antagonist |
| 7 | acetarsol | -0.272 |  |  |  |
| 8 | diroximel-fumarate | -0.267 |  |  | anti-inflammatory agent |
| 9 | pelitinib | -0.262 | 0.850 | EGFR | EGFR inhibitor |
| 10 | resibufogenin | -0.254 |  |  | Na/K-ATPase inhibitor |
| 11 | neratinib | -0.252 | 0.754 | EGFR, ERBB2, KDR | EGFR inhibitor |
| 12 | trichostatin-a | -0.252 | 0.574 | HDAC1, HDAC2, HDAC3, HDAC4, HDAC5, HDAC6, HDAC7, HDAC8, HDAC9, HDAC10, | HDAC inhibitor |
| 13 | cyclovalone | -0.240 | 0.995 | ABCG2 | breast cancer resistance protein inhibitor |
| 14 | NSC-632839 | -0.239 | 0.909 | SENP2, USP1, USP2, USP7 | ubiquitin specific protease inhibitor |
| 15 | LY2874455 | -0.239 | 0.701 | FGFR1, FGFR2, FGFR3, FGFR4, KDR | FGFR antagonist |
| 16 | pacritinib | -0.236 | 0.875 | FLT3, JAK1, JAK2, JAK3 | FLT3 inhibitor, JAK inhibitor |
| 17 | degarelix | -0.228 |  | GNRHR | gonadotropin releasing factor hormone receptor antagonist |
| 18 | nexturastat-a | -0.227 | 1.348 | HDAC1, HDAC6 | HDAC inhibitor |
| 19 | rubitecan | -0.224 | 0.287 | TOP1 | topoisomerase inhibitor |
| 20 | resminostat | -0.222 | 0.944 | HDAC1, HDAC3, HDAC6, HDAC8 | HDAC inhibitor |
